# Gut Microbiome Contributes to Cold-climate Adaptation in Lizards

**DOI:** 10.1101/2022.02.14.480473

**Authors:** Jun-Qiong Chen, Lu-Wen Zhang, Ru-Meng Zhao, Hai-Xia Wu, Long-Hui Lin, Peng Li, Hong Li, Yan-Fu Qu, Xiang Ji

## Abstract

The metabolic cold-climate adaption hypothesis predicts that animals from cold environments have relatively high metabolic rates compared with their warm-climate counterparts. However, studies testing this hypothesis are sparse. Here, we compared gut microbes between two cold-climate lizard species of the genus *Phrynocephalus* to test the hypothesis that gut microbiota can help lizards adapt to cold environments by promoting metabolism and absorption efficiency. We kept lizards at 24°C and 30°C for 25 d, and then collected their fecal samples to analyze and compare the microbiota based on 16S rRNA gene sequencing technology. The gut microbiota was mainly composed of Proteobacteria, Firmicutes, Bacteroidetes, and Verrucomicrobia at the phylum level in both species (Proteobacteria > Firmicutes > Verrucomicrobiota in *P. erythrurus*, and Bacteroidota > Proteobacteria > Firmicutes in *P. przewalskii*). Further analysis revealed that the gut microbiota contributed to the host’s cold adaptation, but with differences in the relative abundance of these contributory bacteria between the two species. KEGG analysis revealed that the gut microbiota primarily played roles in metabolism, genetic information processing, cellular processes, and environmental information processing in both species. Furthermore, genes related to metabolism were more abundant in *P. erythrurus* at 24 °C than in other species × temperature combinations, indicating the role of gut microbiota in long-term cold-climate adaptation. Our finding that gut microbiome contributes to cold-climate adaptation in both species but more evidently in *P. erythrurus* using colder habitats than *P. przewalskii* throughout a year confirms the gut microbiota’s role in the cold-climate adaptation in lizards.

**IMPORTANCE:** This study proves that temperature affects the composition and relative abundance of the gut microbiota in two *Phrynocephalus* lizards in a species-specific manner. Both species harbor specific gut microbiota with significant roles in cold-climate adaptation. Specifically, *P. erythrurus* has a higher Proteobacteria ratio and relative abundance of metabolism-related microbial genes in the gut than *P. przewalskii*. Given that *P. erythrurus* uses colder habitats than *P. przewalskii* throughout a year, these results suggest that gut microbiota contributes to cold-climate adaptation in both species lizards but more evidently in *P. erythrurus*. This study provides evidence linking gut microbiome with cold adaptation. The co-evolution mechanism between gut microbiota and their hosts in extreme environments will provide new insights into animal adaptation.

---

Temperature is a major environmental factor that influences many biological aspects of organisms, including morphology (1), physiology (2), behavior (3), and distribution (4). The metabolic cold adaption hypothesis (MCA) or Krogh’s rule predicts that animals from cold environments have relatively high metabolic rates than those from warmer regions (5, 6). High metabolic rates help animals balance the adverse effects of cold environments by accelerating their physiological processes in a relatively short time (5, 7). Animals adapt to cold environments through physiological (5) and genomic adjustments (8). For example, blackspotted topminnows (*Fundulus olivaceus*) with a northerly (colder) distribution have a higher metabolic rate than conspecifics with a southerly (warmer) distribution (5). Another example is that adaptation to cold environments is the major driving force for evolution of *TRPM8* (a gene codes for a cold-sensing ion channel) in humans (9). However, the role of gut microbiota in cold adaptation is uncertain. Although several studies have demonstrated that temperature significantly influences the gut microbiota and thus host adaptation (10–12), studies on its role in the host’s cold adaptation are rare.

The mutualistic relationship between the host and its gut microbiota has been known for decades. Host animals provide suitable habitats and sufficient nutrients for the survival of gut microorganisms, which, in turn, contribute to nutrient absorption (13), intestinal homeostasis (14), and energy balance (15) of the host by encoding more genes than the host genome (16). Thus, gut microorganisms play a crucial role in host survival (17) and adaptation (18). Temperature affects the composition and relative abundance of host gut microbiota (10–12). Gut microbiota changes at low temperatures are more conducive to the survival and adaptation of host animals in cold climates (10, 14, 19). In mice, the gut microbial changes in cold environments increase the gut absorptive surface area by lengthening the intestinal villi and microvilli, thus enhancing energy extraction (14). In salamanders, cold temperature is correlated with high alpha diversity and improved digestive performance (20).

Reptiles are the first land vertebrates in the amniotic phylogeny and rely on the environment to regulate their body temperate. Does gut microbiota contribute to cold adaptation in reptiles? We already know that gut microbiota has contributed to adaptation to various habitats in reptiles (21–25). For example, it helps invasive red-eared slider turtles (*Trachemys scripta elegans*) colonize new habitats better than native Chinese three-keeled pond turtles (*Chinemys reevesii*) (24). Another example is that the relative abundance of specific bacteria changes with altitude and is associated with adaptation to the Qinghai-Tibet Plateau in Qinghai toad-headed lizards (*Phrynocephalus vlangalii*) (25). Therefore, understanding the coevolution of reptiles and their gut microbiota will reveal their complex environmental adaptation mechanism.

Toad-headed lizards of the reproductively bimodal genus *Phrynocephalus* (Agamidae) have a Palaearctic distribution and occur in desert, arid or semi-arid regions in Central and West Asia and North-Northwest China (26). *Phrynocephalus erythrurus* (viviparous) and *P. przewalskii* (oviparous) studied herein have a strong research foundation in the fields of physiology (2), life history characteristics (27) and molecular biology (28). Both species occur in cold-climate regions, with *P. erythrurus* using colder habitats than *P. przewalskii* throughout a year (Fig. 6). An earlier study on *P. przewalskii* reveals a higher digestive coefficient and assimilation efficiency at low temperature (25 °C) than at medium (33 °C) and high (39 °C) temperatures (2), indicating the potential role of gut microbiota in their cold-climate adaptation. However, no studies have elucidated the contribution of the gut microbiota to cold-climate adaptation in *Phrynocephalus* lizards.

**FIG 6.**
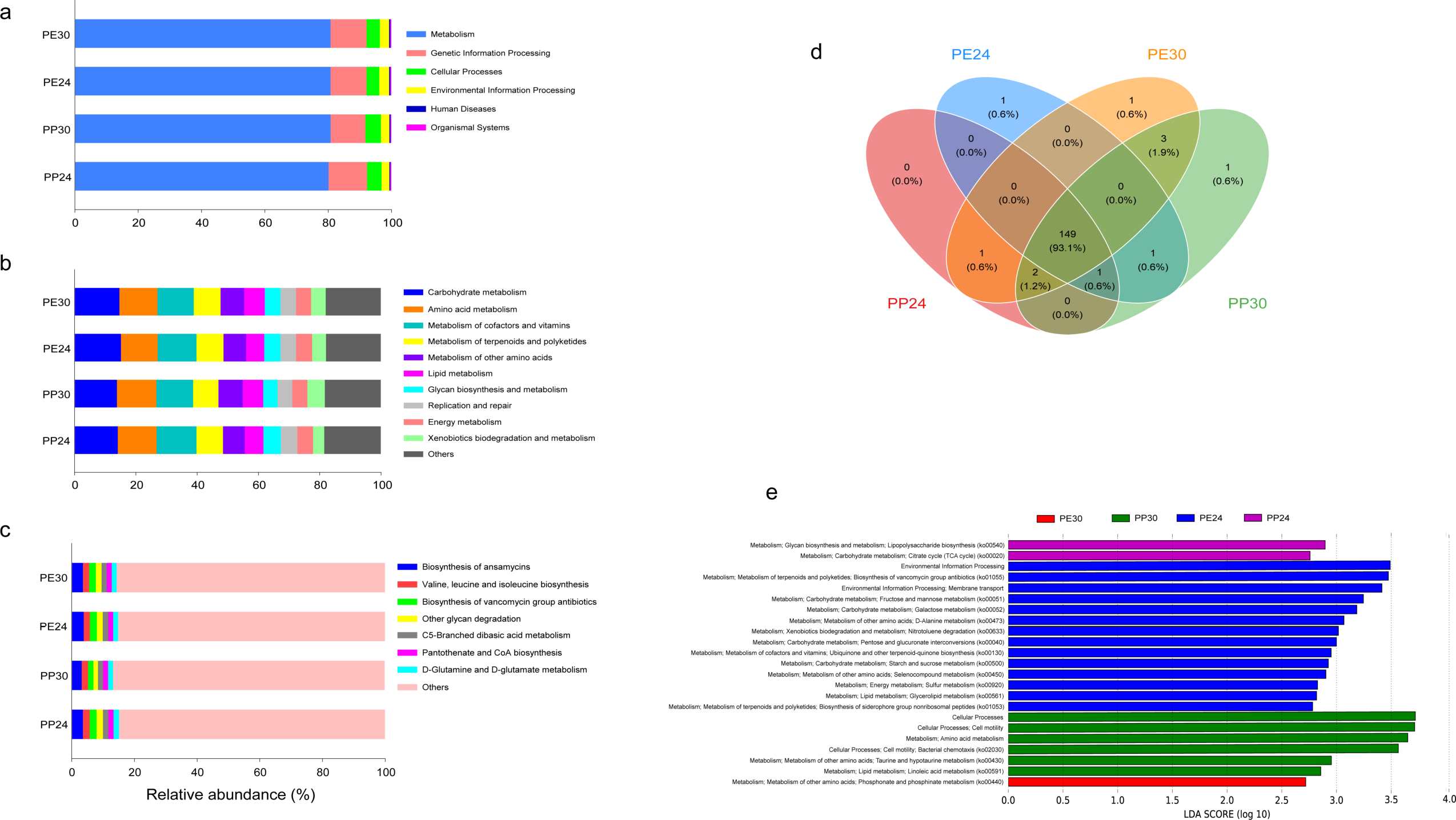
Monthly mean air temperatures and monthly mean rainfall at two localities, from which adult females of *P. erythrurus* (△, Namucuo, Tibet) and *P. przewalskii* (▲, Zhongwei, Ningxia) were collected.

Here, we studied the gut microbiota of *P. erythrurus* and *P. przewalskii* and their role in host adaptability to prolonged cold environments. We analyzed the gut microbiota of these two *Phrynocephalus* lizards maintained at two temperatures (24 °C and 30 °C) to test the following two hypotheses: 1) the gut microbiota can help lizards adapt to cold environments by improving metabolism and absorption efficiency in a species-specific manner, and 2) the gut microbiota from colder environments (*P. erythrurus*) are more helpful to their hosts compared with those from warmer environments (*P. przewalskii*).

## RESULTS

### Fecal bacterial sequencing and AVSs

A total of 1,999,484 and 2,072,977 high-quality reads were obtained from *P. erythrurus* and *P. przewalskii*, respectively, after quality control. SRA accession number and group information are given in supplemental Table S1. The rarefaction curves (Fig. S1) and species accumulation curves (Fig. S2) increased with sample size and attained a precise saturation level, indicating adequate sampling and sufficient sequencing depth in the experiment. Further, based on 99% ASVs full-length sequence reference database, we identified 992 ASVs from the 14 *P. erythrurus* samples (44–265 ASVs per sample) and 1518 ASVs from the 14 *P. przewalskii* samples (147–371 ASVs per sample) (Table S2). Among these ASVs, 1778 from the 56 fecal samples of the two species were taxonomically assigned to 15 phyla, 22 classes, 63 orders, 102 families, and 191 genera.

### Composition and abundance of gut microbiota

We first analyzed the bacterial taxa with high relative abundance in the two species. Proteobacteria (29.79 ± 3.55%), Firmicutes (23.83 ± 1.45%), Bacteroidetes (23.46 ± 1.80%), and Verrucomicrobia (17.05 ± 2.09%) were the most dominant bacterial phyla in both species (Fig. 1a), while Akkermansiaceae (17.05 ± 2.09%), Enterobacteriaceae (10.12 ± 2.45%), Burkholderiaceae (9.18 ± 1.75%), Bacteroidaceae (8.86 ± 0.76%), Caulobacteraceae (6.45 ± 1.34%), Tannerellaceae (5.07 ± 0.62%), and Lachnospiraceae (5.00 ± 0.48%) were the top seven dominant bacterial families (Fig. 1b). *Akkermansia* (17.05 ± 2.09%), *Burkholderia-Caballeronia-Paraburkholderia* (9.17 ± 1.75%), and *Bacteroides* (8.86 ± 0.76%) were the dominant bacterial genera in both species (Fig. 1c).

**FIG 1.**
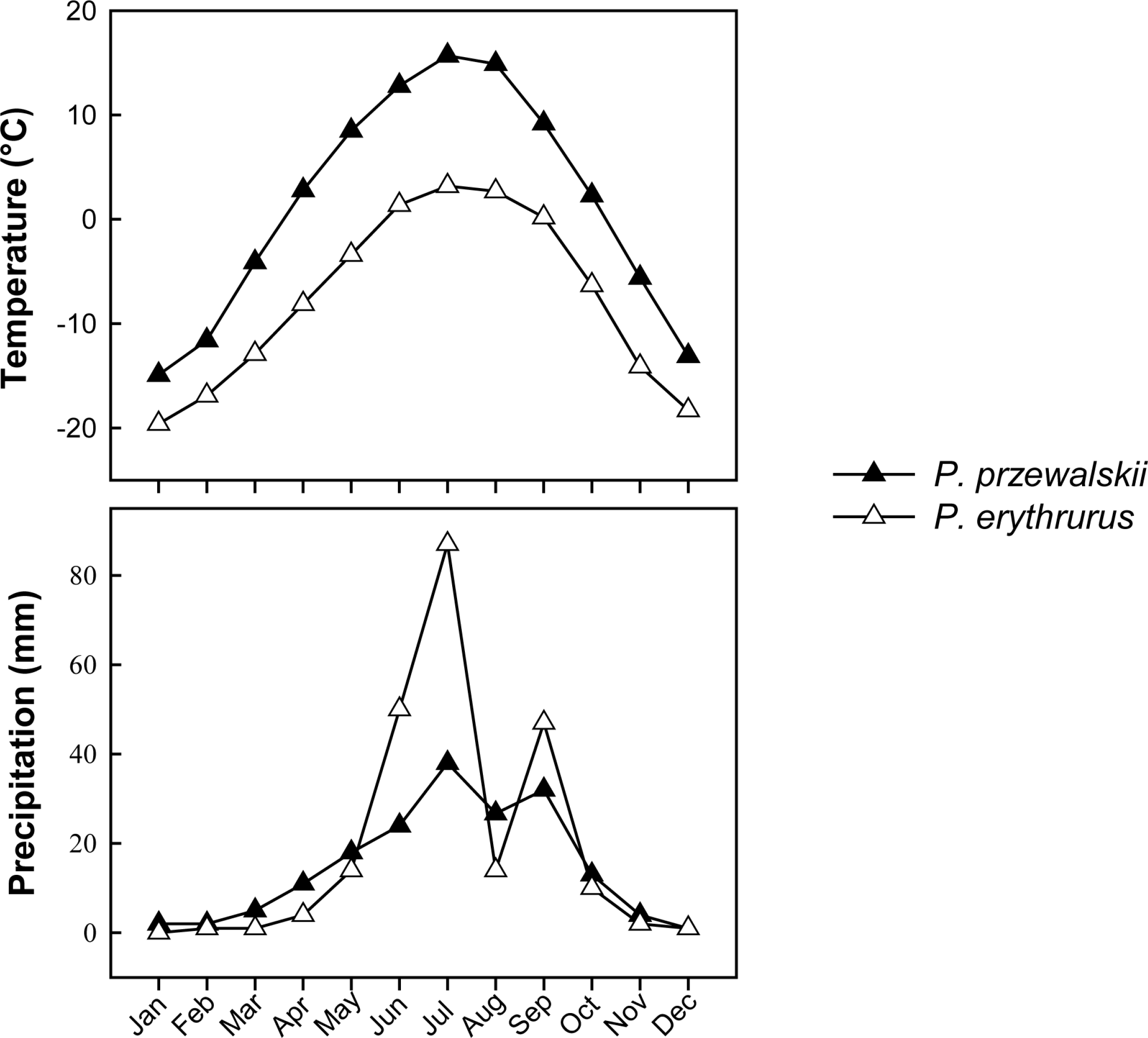
The relative abundance of the gut microbiota at the phylum (a), family (b), and genus (c) levels in four species × temperature combinations. One color indicates one taxon in each plot, and the color for “others” indicates all other taxa not listed in each plot. PP24: *P. przewalskii* at 24 °C; PP30: *P. przewalskii* at 30 °C; PE24: *P. erythrurus* at 24 °C; PE30: *P. erythrurus* at 30 °C.

Additionally, the two lizard species showed significant differences in relative abundance of the different gut microbial taxa. The dominant bacterial phyla in *P. przewalskii* were Bacteroidota (30.84 ± 0.11%) and Proteobacteria (25.46 ± 4.36%), followed by Firmicutes (24.18 ± 1.90%), Verrucomicrobiota (11.81 ± 1.99%), and Desulfobacterota (4.08 ± 0.53%), while those in *P. erythrurus* were Proteobacteria (34.12 ± 5.47%) and Firmicutes (23.47 ± 2.18%), followed by Verrucomicrobiota (22.32 ± 3.39%) and Bacteroidota (16.09 ± 1.90%) (Fig. 1a). Akkermansiaceae (17.05 ± 0.02%) was the most abundant bacterial family in both the lizard species, followed by Burkholderiaceae (11.07 ± 2.46%), Bacteroidaceae (9.52 ± 0.96%), Tannerellaceae (7.60 ± 0.94%), and Caulobacteraceae (7.31 ± 1.95%) in *P. przewalskii* and Enterobacteriaceae (16.61 ± 4.38%), Bacteroidaceae (8.21 ± 1.17%), Burkholderiaceae (7.28 ± 2.43%), and Enterococcaceae (6.12 ± 1.86%) in *P. erythrurus* (Fig. 1b). Meanwhile, *Akkermansia* (17.05 ± 0.02%), *Burkholderia-Caballeronia-Paraburkholderia* (9.17 ± 1.75%), and *Bacteroides* (8.86 ± 0.76%) were the dominant bacterial genera in both the species, followed by *Odoribacter* (4.60 ± 0.80%), *Alistipes* (4.14 ± 0.63%), and *Parabacteroides* (2.72 ± 0.50%) in *P. przewalskii* and *Enterococcus* (3.37 ± 1.67%) in *P. erythrurus* (Fig. 1c).

### Alpha and beta diversity of gut microbiota

Two-way ANOVA showed that Chao1 (*F*_1,52_ = 54.45, *P* < 0.0001) and Shannon (*F*_1,52_ = 36.44, *P* < 0.0001) indexes differed significantly between *P. erythrurus* and *P. przewalskii*; however, Simpson’s eveness index showed no difference (*F*_1,52_ = 0.09 *P* = 0.76). Chao1 and Shannon diversity indexes were higher in *P. przewalskii* than in *P. erythrurus* (Fig. 2). Meanwhile, temperature treatment (*F*_1,52_ < 0.92 and *P* > 0.34) and the interaction between species and temperature (*F*_1,52_ < 2.52 and *P* = 0.11) showed no significant effect on these three indexes. Subsequent PCoA analysis showed a significant separation in gut microbiota among the four species × temperature combinations (ANOSIM: *R* = 0.58, *F*_3,52_ = 6.21, *P* = 0.001; Fig. 3a), with PCo1 and PCo2 accounting for 24% and 14% of the total variance, respectively (Fig. 3a). The PCoA analysis separated the two species (Fig. 3a). Therefore, we performed another PCoA analysis on the two species, respectively. The results showed a significant separation in bacterial composition between the two temperature treatments of each species, with PCo1 and PCo2 together explaining 47% of the total variance in *P. erythrurus* (ANOSIM: *R* = 0.19, *F*_1,26_ = 2.88, *P* = 0.01; Fig. 3b) and 33% of the total variance in *P. przewalskii* (ANOSIM: *R* = 0.18, *F*_1,26_ = 2.17, *P* = 0.01; Fig. 3c).

**FIG 2.**
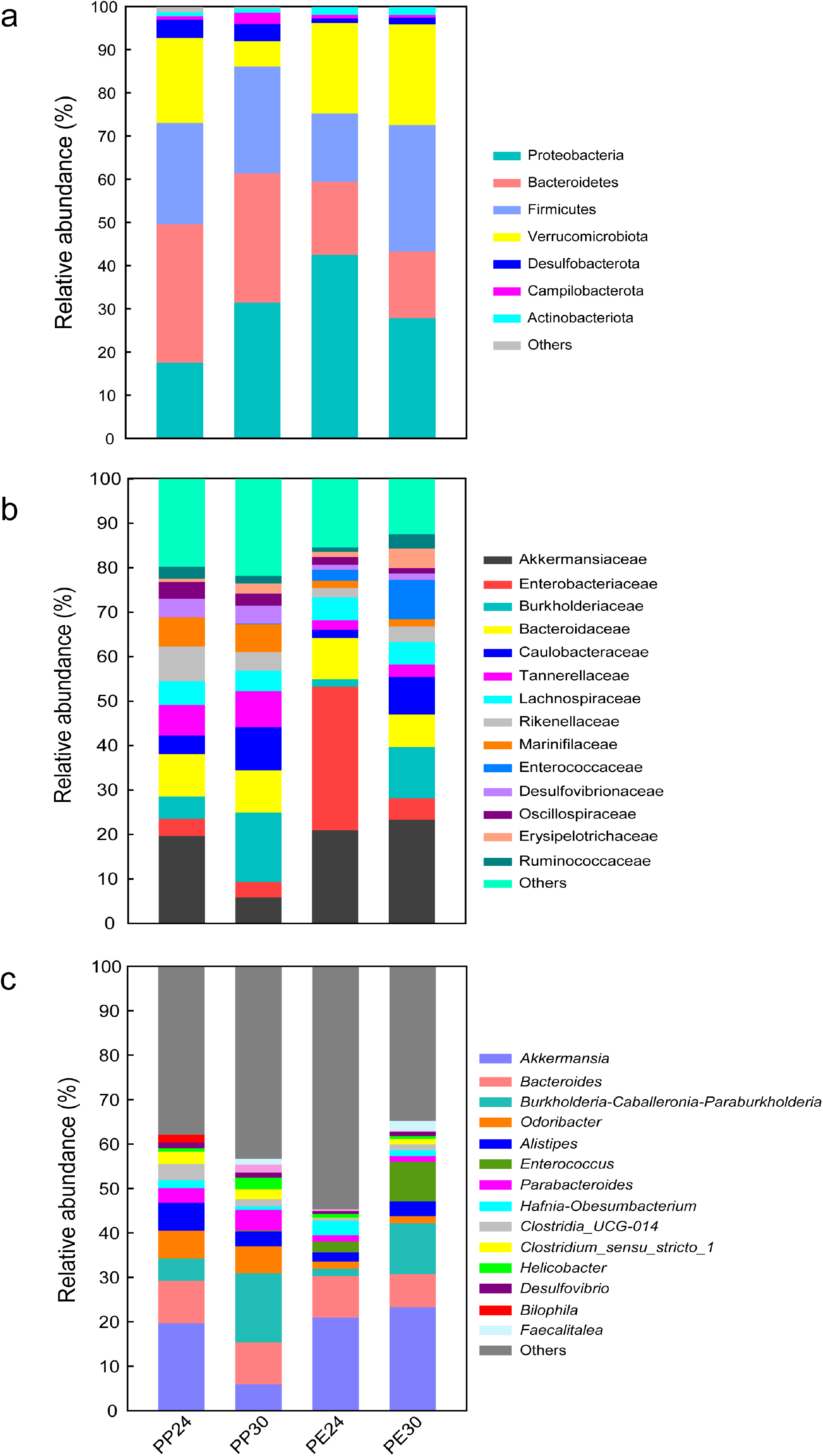
Chao1 index (a), Shannon-Weiner index (b), and Simpson’s eveness index (c) of gut microbiota in four species × temperature combinations. See Figure 1 for the definition of each combination.

**FIG 3.**
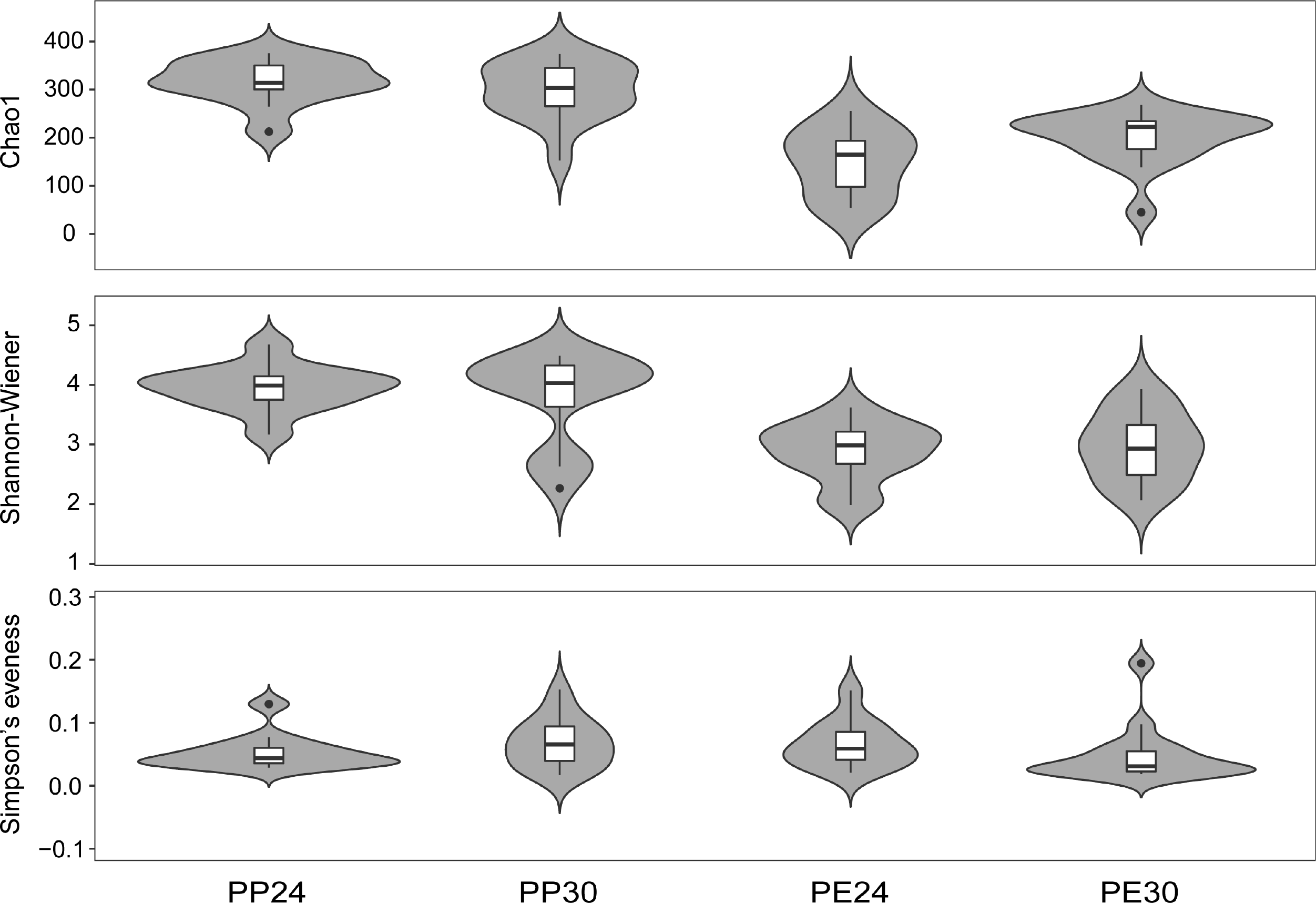
Results of principal coordinates analysis of Bray-Curtis distance matrix for gut microbial diversity in four species × temperature combinations (a), *P. erythrurus* at 24 °C and 30 °C (b), and *P. przewalskii* at 24 °C and 30 °C (c). One color indicates one species × temperature combination. See Figure 1 for the definition of each combination.

Furthermore, the LEfSe and LDA analyses revealed that Enterobacteriaceae (*LDA* = 5.10, *P* = 0.001) was the dominant bacterial family in *P. erythrurus* at 24 °C (Fig. 4), while Erysipelotrichaceae (*LDA* = 4.29, *P* = 0.002) at the family level and *Caproiciproducens* (*LDA* = 3.68, *P* < 0.0001), *Tyzzerella* (*LDA* = 3.58, *P* = 0.04) and *Enterococcus* (*LDA* = 4.62, *P* = 0.0002) at the genus level were dominant at 30 °C (Fig. 4). In *P. przewalskii*, Desulfovibrionaceae (*LDA* = 4.13, *P* = 0.0003) at the family level, and *Alistipes* (*LDA* = 4.30, *P* = 0.03), *Tannerellaceae* (*LDA* = 3.54, *P* = 0.0003), *Rikenella* (*LDA* = 3.55, *P* < 0.0001), *NK4A214 group* (*LDA* = 3.64, *P* = 0.0001), and *Clostridium sensu stricto 1* (*LDA* = 4.10, *P* = 0.0007) at the genus level were dominant at 24 °C (Fig. 4), while *Eubacterium coprostanoligenes group* (*LDA* = 4.21, *P* < 0.0001), *Bacillus* (*LDA* = 3.86, *P* = 0.0003) and *Helicobacter* (*LDA* = 3.98, *P* = 0.004) at the genus level were dominant at 30 °C (Fig. 4).

**FIG 4.**
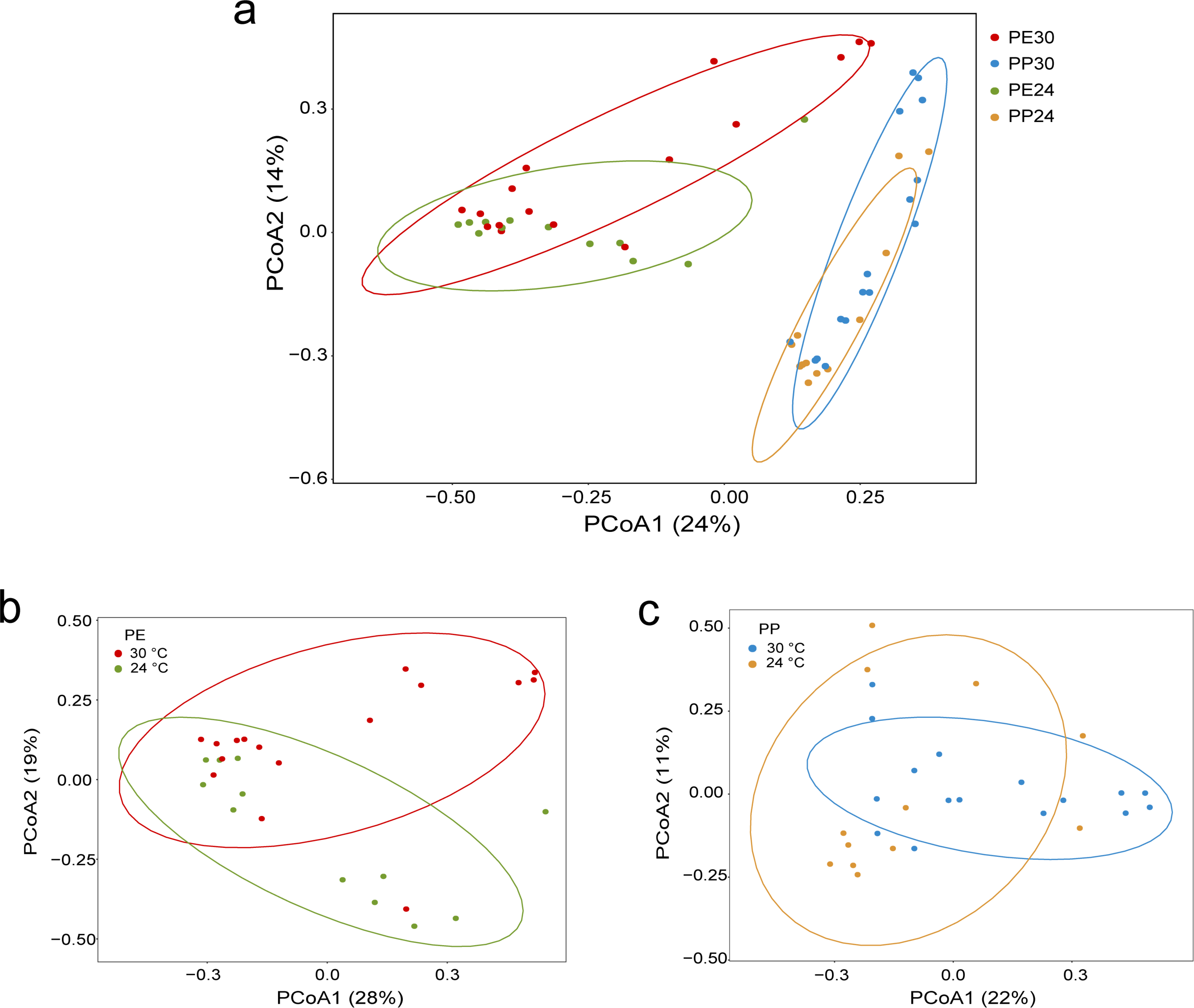
Differences in bacterial taxa among four species × temperature combinations determined by LEfSe (a). LDA scores reflect the differences in relative abundance among four combinations (b). Letters p, c, o, f and g indicate phylum, class, order, family and genus, respectively. See Figure 1 for the definition of each combination.

### The predicted metagenomes

Finally, the gene functions were predicted based on 16S RNA data of the 56 fecal samples. The analysis indicated that genes related to metabolism (80.62 ± 0.12%) were the most predominant at the top level in both species (Fig. 5a). The other three major functional categories at the top level were genetic information processing (11.42 ± 0.20%), cellular processes (4.45 ± 0.10%), and environmental information processing (2.79 ± 0.08%) (Fig. 5a). The metabolism-related functional category included carbohydrate metabolism (14.40 ± 0.12%), amino acid metabolism (12.50 ± 0.08%), metabolism of cofactors and vitamins (12.32 ± 0.15%), metabolism of terpenoids and polyketides (8.60 ± 0.10%), metabolism of other amino acids (7.55 ± 0.12%), lipid metabolism (6.41 ± 0.12%), glycan biosynthesis and metabolism (5.15 ± 0.18%), energy metabolism (5.04 ± 0.04%), and xenobiotics biodegradation and metabolism (4.74 ± 0.31%) as the predominant ones at the second level in both species (Fig. 5b). Other functional categories with a relative abundance greater than 3% included replication and repair (5.07 ± 0.10%) at the second level and genetic information processing at the first level in both species (Fig. 5b). At the third level, the most abundant gene functions in the four lizard groups were biosynthesis of ansamycins (3.54 ± 0.12%), valine, leucine, and isoleucine biosynthesis (2.04 ± 0.02%), and biosynthesis of vancomycin group antibiotics (2.03 ± 0.07%) (Fig. 5c).

**FIG 5.**
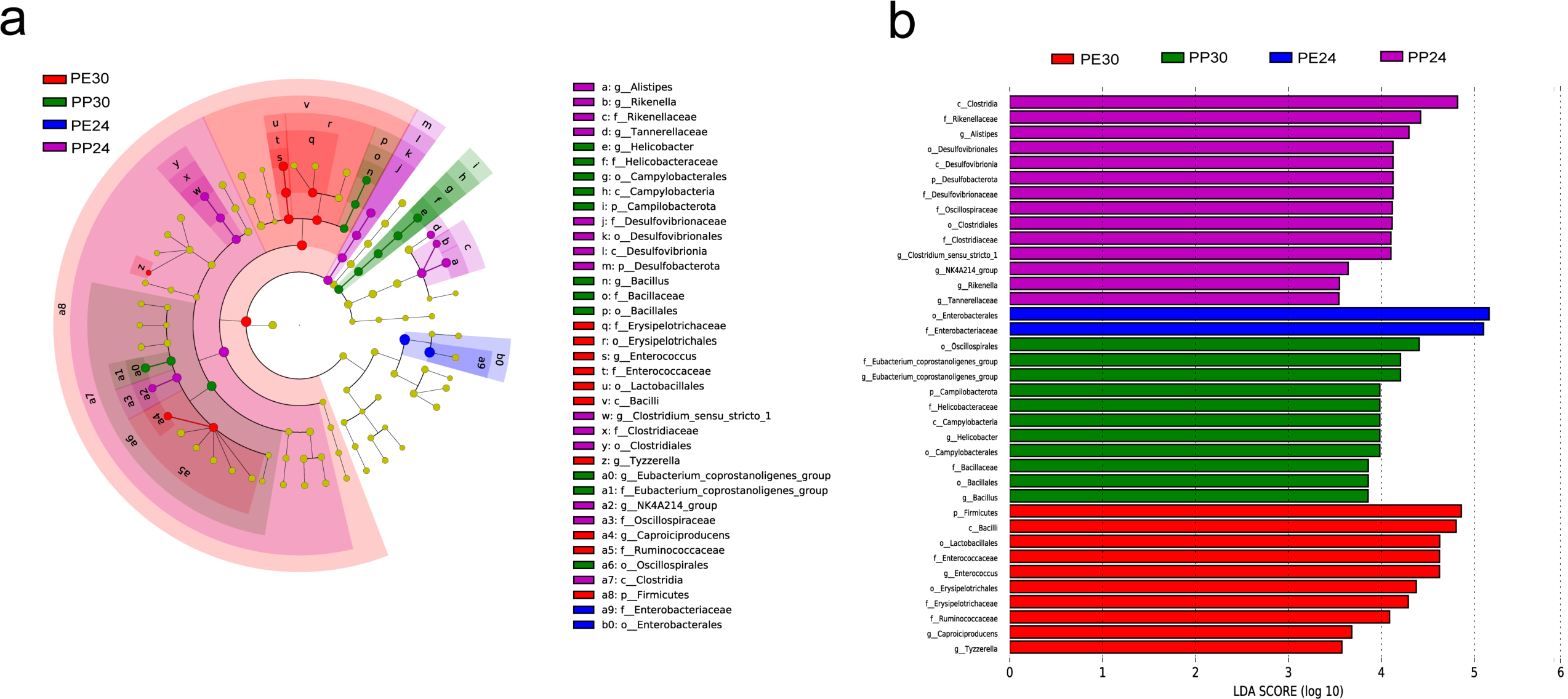
Relative abundance of gene functions at the first (a), second (b), and third (c) levels in the gut microbiota of two lizard species, and the Venn diagram showing the functional genes among fecal samples from four species × temperature combinations (d). LDA scores reflect the differences in relative abundance among the four groups of fecal samples (e). One color indicates one gene function in each plot, wiht detailed descriptions shown on the right side of each plot. The colors for “others” in figures b and c indicate all other gene functions not listed in these two plots. See Figure 1 for the definition of each combination.

Furthermore, 160 known KEGG genes were identified based on 16S rRNA sequences in the two species. Venn diagram showed that most of the KOs were common in the different species × temperature combinations (Fig. 5d); 154 KOs were shared between the two species, and 149 between the 24 °C and 30 °C treatments in *P. erythrurus*, and 152 between the two temperature treatments in *P. przewalskii*. The gene functions enriched at 24 °C in *P. erythrurus* were related to metabolism, including phosphonate and phosphinate metabolism (ko00440; *LDA* = 3.00, *P* = 0.0009), fructose and mannose metabolism (ko00051; *LDA* = 3.25, *P* = 0.001), galactose metabolism (ko00052; *LDA* = 3.18, *P* = 0.02), starch and sucrose metabolism (ko00500; *LDA* = 2.92, *P* = 0.03), sulfur metabolism (ko00920; *LDA* = 2.83, *P* = 0.0006), glycerolipid metabolism (ko00561; *LDA* = 2.82, *P* = 0.002), ubiquinone and other terpenoid-quinone biosynthesis (ko00130; *LDA* = 2.95, *P* = 0.0009), selenocompound metabolism (ko00450; *LDA* = 2.90, *P* = 0.0006), D-alanine metabolism (ko00473; *LDA* = 3.07, *P* = 0.001), biosynthesis of siderophore group nonribosomal peptides (ko01053; *LDA* = 2.78, *P* = 0.0001), biosynthesis of vancomycin group antibiotics (ko01055; *LDA* = 3.47, *P* = 0.02), and nitrotoluene degradation (ko00633; *LDA* = 3.02, *P* = 0.001), and environmental information processing, including membrane transport (*LDA* = 3.41, *P* = 0.009) (Fig. 5e). However, only ko00440 related to phosphonate and phosphinate metabolism (*LDA* = 2.72, *P* = 0.02) showed a high relative abundance at 30 °C in *P. erythrurus* (Fig. 5e). Gene function related to metabolism, including TCA cycle (ko00020; *LDA* = 2.76, *P* = 0.01) and lipopolysaccharide biosynthesis (ko00540; *LDA* = 2.90, *P* = 0.007), were enriched at 24 °C in *P. przewalskii* (Fig. 5e). Meanwhile, *P. przewalskii* at 30 °C owned relatively more KOs related to cellular processes, including bacterial chemotaxis (ko02030; *LDA* = 3.57, *P* = 0.006), and metabolism-related functions, including amino acid metabolism (*LDA* = 3.65, *P* = 0.0003), linoleic acid metabolism (ko00591; *LDA* = 2.86, *P* = 0.003), and taurine and hypotaurine metabolism (ko00430; *LDA* = 2.95, *P* = 0.002) (Fig. 5e). In addition, *P. erythrurus* had higher abundance of gene functions related to metabolism at 24 °C than *P. przewalskii*.

## DISCUSSION

Investigating the differences in gut microbiota associated with cold-climate and/or high-altitude adaptation can contribute to uncovering the mechanisms of cold adaptation in animals. In this study, the alpha diversity of gut bacterial communities differed between the two species but not between the two temperature treatments (Fig. 2). Similarly, temperature does not significantly affect the gut bacterial alpha-diversity in cows (29) or tadpoles (30). However, most studies spanning a wide range of animal taxa, including invertebrates and vertebrates (12, 14, 20−21, 31), have shown that the gut bacterial alpha-diversity overall increases at high temperatures; the opposite has been observed in fish (10). Past studies generally conclude that the gut microbial diversity in animals is related to their genotype or habitat environment (22, 25).

Both species studied herein live in cold and unstable environments. Therefore, the gut microbiota might have adapted to drastic temperature fluctuations during evolution (20). In the present study, the alpha diversity was significantly higher in *P. przewalskii* than in *P. erythrurus*. This observation is consistent with the study on the Qinghai toad-headed lizard *P. vlangalii*, where the lowest gut microbial diversity is detected in the highest altitude (coldest) population (25). However, these results are different from those reported for mammals. High-altitude mammals (e.g., yaks, sheep, and pika) have higher bacterial diversity than their low-altitude relatives (cattle, sheep, and pika) due to their higher energy demands in cold and hypoxic environments (32, 33); however, the cause for these differences between reptiles and mammals remains unknown. In addition, many studies have shown that the host genotype causes differences in the alpha-diversity in different host taxa (13, 33−34). For example, the alpha diversity of gut microbiota differs significantly among four species of fish larvae reared under the same conditions, indicating the influence of the genetic background of a species on the alpha diversity (34). Therefore, the differences in gut microbial diversity between the two *Phrynocephalus* lizards observed in this study suggest an association with their genetic background.

The gut microbiota plays a vital role in animal adaptation and evolution (22, 25). Past studies have reported significant differences in the composition of gut microbes at different taxon levels in reptiles due to various factors, including host genotype (35), dietary type (36), habitat use (25), gender (37), and ontogenetic stage (30). The present study also found significant differences in the relative abundance of gut bacteria at different taxon levels among the different species and temperature combinations (Fig. 3 and 4). The top two dominant phyla of gut microbiomes in *P. erythrurus* were Proteobacteria and Firmicutes (Fig. 1), consistent with *P. vlangalii* (25) and the marine iguana *Amblyrhynchus cristatus* (22). Proteobacteria degrades and ferments complex sugars and synthesizes vitamins to supply nutrients for their hosts (38).

Meanwhile, the Firmicutes phylum is associated with enzymes involved in fermentation and vitamin B biosynthesis (39). Therefore, the higher relative abundance of Proteobacteria and Firmicutes in *P. erythrurus* indicates their roles in improving the host energy absorption efficiency and utilization rate. Meanwhile, Bacteroidota and Proteobacteria were the dominant gut bacteria at the phylum level in *P. przewalskii* (Fig. 1), similar to the crocodile lizard *Shinisaurus crocodilurus* (40). Unlike *P. erythrurus*, *P. przewalskii* had more Bacteroidota, correlated with the breakdown of the complex macromolecules (38).

Researchers have also positively correlated the Proteobacteria ratio (the abundance of Proteobacteria divided by the abundance of Firmicutes and Bacteroidetes) with the bacterial stress tolerance under cold environments (41). In this study, *P. erythrurus* (0.86) had a higher Proteobacteria ratio than *P. przewalskii* (0.46), which indicates that *P. erythrurus* harbor gut microbiota that helps the host’s adapt to cold environments compared with those found in *P. przewalskii* in warmer climates. Comparing the difference in values between the two temperature treatments, we found that *P. erythrurus* had the largest Proteobacteria ratio (1.30) at 24 °C, followed by *P. erythrurus* at 30 °C (0.62) and *P. przewalskii* (0.31 at 24 °C and 0.58 at 30 °C). This suggests that the Proteobacteria ratio contributes to the cold adaptation of animals, and there are also inter-specific differences. In addition, we found that the ratio of Firmicutes to Bacteroidetes was arranged in order of *P. przewalskii* at 24 °C < *P. przewalskii* at 30 °C < *P. erythrurus* at 24 °C < *P. erythrurus* at 30 °C. The ratio of Firmicutes to Bacteroidetes is negatively correlated with mass gain (42). The Firmicutes/Bacteroidetes ratio suggests that the gut microbiota could rapidly increase the host mass at low temperatures, which is conducive to the host’s adaptation to the long-term cold environment.

Several studies have proven the contribution of gut microbiota to the host’s adaptation to cold environments (10, 14, 19). An increase in the relative abundance of gut bacterial members that can degrade host-derived mucin glycans helps Chinese alligators cope with cold climates (43). However, the contribution of gut microbiota to cold adaptation is not the same across animals (10). In the 24 °C treatment, bacteria of the family Enterobacteriaceae were enriched in *P. erythrurus*, while Desulfovibrionaceae, *Alistipes*, *Tannerellaceae*, *Rikenella*, *NK4A214 group*, and *Clostridium sensu stricto 1* were enriched in *P. przewalskii* (Fig. 4). These bacteria, correlated with fermentation and metabolism and relatively abundant at 24 °C in both the species, probably contributed to the host’s adaptation to cold environments. Generally, bacteria of the family Enterobacteriaceae are associated with glucose fermentation and reduces nitrates to nitrites (44), whereas Desulfovibrionaceae bacteria produce hydrogen sulfide through sulfate reduction (45), which increases the mucus layer permeability (46). Bacteria of the genera *Alistipes* and *Rikenella* are correlated with metabolism (47). Meanwhile, *Clostridium sensu stricto 1* accelerates the use of cellulose in the host (48). In conclusion, significant differences detected in these bacteria suggest their contribution to cold-climate adaptation in these two *Phrynocephalus* lizards.

Similarly, the two species had different dominant bacteria at 30 °C (Fig. 1 and 4). Bacteria of the family Erysipelotrichaceae and genera *Caproiciproducens*, *Tyzzerella*, and *Enterococcus* were more in *P. erythrurus*, and those of the genera *Bacillus*, *Helicobacter* and the *Eubacterium coprostanoligenes group* were more in *P. przewalskii* (Fig. 1 and 4). These bacteria, relatively abundant at 30 °C in both species, play metabolism-related roles during digestion and absorption, but their specific functions differed from 24 °C. For example, the family Erysipelotrichaceae is probably related to the host’s lipid metabolism and contributes to host nutrient absorption in *P. erythrurus* (49). Bacteria of the genus *Caproiciproducens* increase caproic acid production (50), while bacteria of the genus *Enterococcus* enhance anticoagulation capacity (51). Meanwhile, the *Eubacterium coprostanoligenes group* family in *P. przewalskii* is correlated with the host lipid homeostasis (52). These results collectively indicate that the two *Phrynocephalus* lizards have had a more tolerant gut microbiome than other animals. This mechanism contrasts with other known species, where the gut microbiota in high-temperature environments is composed of mostly microbes harmful to their hosts (12, 23). For example, the warm-adapted genus *Mycobacterium* exhibits higher abundances in *Rana pipiens* tadpoles reared at warm temperatures (30). Meanwhile, the heat-induced changes in the gut microbiota in laying hens are associated with hepatic and intestinal dysfunction (12).

Furthermore, the gene functions predicted based on the 16S rRNA sequences of gut microbiota in the two *Phrynocephalus* lizards were mainly metabolism-related, consistent with other known lizards (36, 53). Comparison of the two *Phrynocephalus* lizards at different temperatures revealed significant differences in the functions of the gut microbiota among different species and temperature combinations (Fig. 5). The gut microbes in *P. erythrurus* at 24 °C were primarily involved in environmental information processing and metabolism (including 12 KOs were mainly correlated with carbohydrate and salt metabolism) (Fig. 5). In contrast, the most significant gut microbial function in *P. przewalskii* at 24 °C, including two KOs, was related to only metabolism (Fig. 5). In addition, significant differences were detected between the two species in gut microbial function at 30 °C. The only KO related to metabolism was prominent at 30 °C in *P. erythrurus*, while the most significant function at 30 °C in *P. przewalskii* was related to amino acid metabolism (including three KOs) and cellular processes (including one KO) (Fig. 5).

*Phrynocephalus erythrurus* at 24 °C had the highest relative abundance in the gut bacterial KOs than other species and temperature combinations, suggesting gut microbiota assists in long-term cold adaptation of *P. erythrurus* through high expression of metabolism-related genes. Above results indicate that gut microbiota from colder environments (*P. erythrurus*) had a higher proportion of metabolism-related functions than those from warmer environments (*P. przewalskii*), which might be more conducive to the host’s cold-climate adaptation.

## MATERIAL AND METHODS

### Animal collection and maintenance

In late July 2020, we collected 14 adult female *P. erythrurus* from Namucuo (30°69’N, 90°87’E), Tibet, and 14 adult female *P. przewalskii* from Zhongwei (37°46’N, 104°87’E), Ningxia. Both places are dry and cold, with Namucuo being colder than Zhongwei throughout a year (Fig. 6) and, as such, oviparous lizards cannot live in the former place (54). The 28 female lizards were brought to our laboratory at Nanjing Normal University, where 4−5 lizards of the same species were housed together in a plastic cage [500 × 400 × 360 mm (length × width × height)] with a sand substrate (∼150 mm depth) and pieces of ceramic tiles for shade and shelter. These cages were placed in a room at temperatures varying from 24−30 °C. Thermoregulatory opportunities were provided during daytime (07:00−19:00 h) using a 60 W incandescent lamp suspend. Our experimental procedures were approved by the Institutional Animal Care and Use Committee of Nanjing Normal University, and were conducted in accordance with related guidelines (IACUC20200511).

### Experiment design and sample collection

We conducted a two temperatures (24 °C and 30 °C) × two species (*P. erythrurus* and *P. przewalskii*) factorial design experiment in September 2020. Before the experiment, two AAPS (artificial atmospheric phenomena simulator) rooms and all experimental items were disinfected with alcohol wipes and an ultraviolet lamp. We individually housed lizards in 350 × 240 × 190 mm plastic cages, and them placed 12 lizards (six of each species) in an AAPS room at 24 ± 1 °C and 16 (eight of each species) in an AAPS room at 30 ± 1 °C. Fluorescent lights in both rooms were switched on at 07:00 h and off at 19:00 h throughout the experiment. Mean values for body mass and snout-vent length did not differ between the two temperatures in both species (*F*_1, 12_ < 0.71 and *P* > 0.41 in all cases). Mealworms (larvae of *Tenebrio molito*r sterilized with an ultraviolet lamp 1 h before feeding) and distilled water were provided *ad libitum* for 25 d. Feces of each lizard were collected in sterile centrifuge tubes under sterile conditions at two intervals, one between day 9 and day 12 and another between day 21 and day 25. All 56 fecal samples were individually labeled and stored at −80 °C for DNA extraction.

### DNA extraction, PCR amplification and sequencing of fecal samples

The total DNA was extracted from the fecal samples using the E.Z.N.ATM Mag-Bind Soil DNA Kit (OMEGA Bio-Tek, Norcross, GA, USA) according to the manufacturer’s instructions. The DNA quality was assessed on an agarose gel (1.0%), and the concentration was detected using a Qubit3.0 DNA detection kit (Thermo Fisher Scientific, Shanghai, China). The V3-V4 region of the bacterial 16S rRNA gene was amplified by polymerase chain reaction (PCR) using the gene-specific primers 341F (5’-CCTACGGGNGGCWGCAG-3’) and 805R (5’-GACTACHVGGGTATCTAATCC-3’). The first round of PCR was conducted in a 30 μL reaction mixture consisting of 15 μL of Hieff Robust PCR Master Mix (2×), 1 μL of each primer (10 μM), 10−20 ng of genomic DNA, and 9−12 μL of ddH_2_O. The thermal cycling conditions were set as follows: initial denaturation at 94 °C for 3 min, followed by 5 cycles of denaturation at 94 °C for 30 s, annealing at 45 °C for 20 s and extension at 65 °C for 30 s, 20 cycles of denaturation at 94 °C for 20 s, annealing at 55 °C for 20 s, and extension at 72 °C for 30 s, and 5 min at 72 °C. The second round of PCR was conducted using the first-round PCR amplicon as the template. The six-base recognition sequence (barcodes) and Illumina bridge PCR compatible primers were introduced in the second PCR amplification after the first. The PCR mixture used was the same as the first round. The thermal cycling conditions were set as follows: denaturation at 95 °C for 3 min, followed by 5 cycles of denaturation at 94 °C for 20 sec, annealing at 55 °C for 20 sec and an extension at 72 °C for 30 sec, and a final extension at 72 °C for 5 min. The PCR products electrophoresed on a 2.0% agarose gel were purified, and the concentration was detected using a Qubit3.0 DNA detection kit (Thermo Fisher Scientific). Subsequently, 10 ng of the PCR product (600 pmol) was used from each sample for pyrosequencing on an Illumina NovaSeq 6000 system at Shanghai Sangon Biotech., China.

### Data process and standardization

We used the Quantitative Insights Into Microbial Ecology 2 (QIIME2) platform (55) to process the raw sequencing reads. Subsequently, we used the DADA2 package (56) to filter and trim the low-quality reads to produce paired-end reads, and merged these paired-end reads to obtain the amplicon sequence variants (ASVs). The taxonomy was assigned to the ASVs using the pre-formatted SILVA 138 SUURef NR99 ASVs full-length reference sequences following the q2-fragment-classifier method in QIIME2. ASVs with the number of ASVs greater than 10 in at least two samples were retained for further analysis to avoid the effect of low read numbers on the results using QIIME2. Finally, the ASV abundance was standardized based on the sample with the least ASVs for alpha and beta diversity analysis.

### Comparison and diversity of gut microbiota

The alpha diversity matrices, including community richness (Chao1 index), community diversity (Shannon diversity index), and community evenness (Simpson’s evenness index), were calculated in QIIME2. We used the R software to visualize these indices. Further, we performed principal coordinates analysis (PCoA) and analysis of similarity (ANOSIM) to determine the differences in the structure (beta diversity) and communities of the gut microbiota among the different species × temperature combinations. The cluster analysis was performed using Bray-Curtis similarity to explore the similarities in the gut community composition between the samples. ANOSIM based on the Bray-Curtis distance metrics with 999 permutations was further conducted to analyze the differences across the groups.

Subsequently, we used linear discriminant analysis effect size (LEfSe) (57) to compare the microbial abundances, from the phylum to family levels, between the two species and between the two temperature treatments. Then, we used linear discriminatory analysis (LDA) to evaluate the effect size for each selected classification. Here, we only used the dominant bacterial taxa with a log LDA score > 3.5 (over 3.5 orders of magnitude).

### Gene function prediction

Further, the gene functions based on 16S rRNA sequences were predicted using PICRUSt2 based on the Kyoto Encyclopedia of Genes and Genomes (KEGG) database (58), and the genes were classified and allocated to the corresponding KEGG pathways (59). Functional genes in each pathway were counted to assess their relative abundance in each species × temperature combination. Further, we used LEfSe to compare the relative abundance of the KEGG gene functions from level 1 to level 3 KOs, thereby determining the differences in functional genes among the different species × temperature combinations. We performed LDA analysis to assess the effect size for each selected level. Here, we only used the functional category with a log LDA score > 2.5.

### Statistical analysis

Neither alpha (*F*_1,10_ < 4.59, *P* > 0.05) nor beta (ANOSIM: *R* < 0.14, *P* > 0.14) diversity differed between samples collected at the two time intervals (Day 9−12 and Day 21−24) within each species × temperature combination. Therefore, we pooled the data for the two intervals. We used two-way analysis of variance (ANOVA) to test the differences in alpha diversity and the relative abundance of the gene functions between the two temperature treatments and between the two species. All values were presented as mean ± standard error (SE), and the differences were considered statistically significant at α < 0.05.

### Data availability

The sequencing dataset for gut microbes has been submitted to the National Genomics Data Center (NGDC) GSA database (accession number CRA004548; https://ngdc.cncb.ac.cn/gsa/s/F0ZwqzKc).

## ACKNOWLEDGMENTS

We thank Kun Guo, Xiang-Mo Li, Xia-Qiu Tao, Fan Xie, and Lin Zhu for help in sample collection during the research. The study was supported by the grants from the National Natural Science Foundation of China (31670422, 31672277, 31870390, and 31971414), the Second Tibetan Plateau Scientific Expedition and Research Program (STEP) (2019QZKK05010216), Jiangsu Provincial Natural Science Foundation (BK20161556), Natural Science Foundation of the Jiangsu Higher Education Institutions (19KJA330001), and Postgraduate Research & Practice Innovation Program of Jiangsu Province (KYCX20_1241).

We declare no conflict of interest.

J.-Q.C., Y.-F.Q., X.J. conceived and designed the experiment; Y.-F.Q., X.J. supervised the study; J.-Q.C., R.-M.Z., L.-W.Z., H.-X.X., H.-X.W., L.-H.L., P.L., H.L., Y.-F.Q. conducted the experiment and collected the data; J.-Q.C., R.-M.Z., L.-W.Z., Y.-F.Q. analyzed the data and conducted data visualization; Y.-F.Q., X.J. wrote the paper with improvement suggestions from all co-authors. All authors reviewed and approved the final manuscript.

